# Deep Brain Stimulation: Imaging on a group level

**DOI:** 10.1101/2020.01.14.904615

**Authors:** Svenja Treu, Bryan Strange, Simon Oxenford, Andrea Kühn, Ningfei Li, Andreas Horn

## Abstract

Deep Brain Stimulation (DBS) is an established treatment option for movement disorders and is investigated to treat a growing number of other brain disorders. It has been shown that DBS effects are highly dependent on exact electrode placement, which is especially important when probing novel indications or stereotactic targets. Thus, considering precise electrode placement is crucial when investigating efficacy of DBS targets. To measure clinical improvement as a function of electrode placement, neuroscientific methodology and specialized software tools are needed. Such tools should have the goal to make electrode placement comparable across patients and DBS centers, and include statistical analysis options to validate and define optimal targets. Moreover, to allow for comparability across different research sites, these need to be performed within an algorithmically and anatomically standardized and openly available group space. With the publication of Lead-DBS software in 2014, an open-source tool was introduced that allowed for precise electrode reconstructions based on pre- and postoperative neuroimaging data. Here, we introduce *Lead Group*, implemented within the Lead-DBS environment and specifically designed to meet aforementioned demands. In the present article, we showcase the various processing streams of *Lead Group* in a retrospective cohort of 51 patients suffering from Parkinson’s disease, who were implanted with DBS electrodes to the subthalamic nucleus (STN). Specifically, we demonstrate various ways to visualize placement of all electrodes in the group and map clinical improvement values to subcortical space. We do so by using active coordinates and volumes of tissue activated, showing converging evidence of an optimal DBS target in the dorsolateral STN. Second, we relate DBS outcome to the impact of each electrode on local structures by measuring overlap of stimulation volumes with the STN. Finally, we explore the software functions for connectomic mapping, which may be used to relate DBS outcomes to connectivity estimates with remote brain areas. We isolate a specific fiber bundle – which structurally resembles the hyperdirect pathway – that is associated with good clinical outcome in the cohort. The manuscript is accompanied by a walkthrough tutorial through which users are able to reproduce all main results presented in the present manuscript. All data and code needed to reproduce results are openly available.

**Highlights:**

- We present a novel toolbox to carry out DBS imaging analyses on a group-level
- Group electrodes are visualized in 2D and 3D and related to clinical regressors
- A favorable target and connectivity profiles for the treatment of PD are validated

## Introduction

The modulation of neural networks by Deep-Brain Stimulation (DBS) is an efficacious and established treatment option for specific neurological and psychiatric disorders, and is currently investigated for other brain disorders. DBS treatment is best explored in movement disorders (Benabid et al., 1991; Kupsch et al., 2006a), with the subthalamic nucleus (STN) and internal pallidum (GPi) as the most established targets (Schuepbach et al., 2013). The application is continuously extended to other indications and targets (for a review see Lozano and Lipsman, 2013). Aside from its clinical value, DBS opens an invaluable window to the human brain and it is increasingly adopted to probe causal relationships between stimulated targets and certain behavioral effects, such as risk control (Irmen et al., 2019a; Nachev et al., 2015), movement speed (W.-J. Neumann et al., 2018), memory learning (de Almeida Marcelino et al., 2019) or verbal fluency (Ehlen et al., 2017; Mikos et al., 2011).

In early clinical studies which led to FDA- and CE-approval for indications like Parkinson’s Disease, Dystonia or Essential Tremor, the exact electrode placements were not investigated (Deuschl et al., 2006; Kupsch et al., 2006b; Schuepbach et al., 2013). However, in these, DBS targets had already been well informed and established by decades of ablative surgery, which leads to about equal clinical effects (Altinel et al., 2019; Starr et al., 1998). However, a multitude of studies have shown that DBS electrodes need to be accurately placed to maximize clinical improvements (e.g. Dembek et al., 2019a; Horn, 2019; Horn et al., 2019a). Thus, when probing novel targets, things could turn out differently. For instance, in Depression and Alzheimer’s Disease, clinicians report that some patients largely improve while others do not (Laxton et al., 2010a; Riva-Posse et al., 2017). In the case of depression, however, two prospective, randomized, sham-controlled trials have failed (Dougherty et al., 2015; Holtzheimer et al., 2012). While some patients had responded well to treatment, their group effects were diminished by non-responders. A possible (but unknown) explanation for the variability in outcome would be that responders had optimal lead placement while non-responders did not. By investigating their differential electrode placements, one could possibly find relationships between clinical outcome and modulated anatomical space and brain connectivity measures, as was shown for other diseases (Al-Fatly et al., 2019; Baldermann et al., 2019; Dembek et al., 2019a; Horn, 2019; Reich et al., 2019).

Thus, large-scale group studies that include electrode placement could comprehensively help to investigate relationships between stimulation sites and clinical/behavioral outcomes.

With the publication of the software Lead-DBS in 2014 (Horn and Kühn, 2015) and its further methodological advancement over the years (Ewert et al., 2019, 2018; Horn et al., 2019a, 2014; Horn and Blankenburg, 2016), an openly available tool was created, which allows for precise electrode reconstructions based on pre- and post-operative imaging data.

One key goal of the software is to make electrode placement transferable and comparable across centers and patients by warping their coordinates into a common stereotactic space. While the idea of warping to MNI space is surely not novel (the concept has been around in the neuroimaging community for decades, e.g. (Ashburner, 2012)), the process still has its limitations in an inherent loss of precision. Errors of placements in DBS electrode locations are prone to occur when studying them in a common stereotactic space. Over the years, our group and others have focused on minimizing these kinds of errors and improving the accuracy of Lead-DBS (Ewert et al., 2019b, 2018b; Hellerbach et al., 2018; Horn et al., 2019a; Husch et al., 2018; Schönecker et al., 2009; Dembek et al., 2019b). Strategies such as multispectral warps, subcortical refinements, brain shift correction, phantom-validated automatic electrode localization, the possibility of manual refinements of warfields and detection of electrode orientation have led to a freely available pipeline that aims at maximizing precision both in native and common space (for an overview see (Horn et al., 2019a)). The registration pipeline of Lead-DBS was recently evaluated in a large comparative study and results were comparable to manual expert delineations of subcortical nuclei (Ewert et al., 2019b). While Lead-DBS is capable to register patient data to different stereotactical spaces, a worldwide standard of the neuroimaging community has been adopted as default: The Montreal Neurological Institute (MNI) space in its most current and best-resolved version (ICBM 2009b Nonlinear Asymmetric space).

Various publications could demonstrate that electrode reconstructions in such common spaces remain informative and may be used to predict outcomes or behavioral measures across cohorts and DBS centers (Al-Fatly et al., 2019; Baldermann et al., 2019; de Almeida Marcelino et al., 2019; Horn et al., 2019a; W.-J. Neumann et al., 2018). Once electrodes are in such a common space, this allows for analyses of DBS electrodes on a group level, and direct comparability of results between patients, research groups and software tools.

Here, we introduce a novel open source toolbox, *Lead Group*, which was implemented within the Lead-DBS environment and specifically designed with group level analyses in mind. While *Lead Group* has been available in prototype form for a while, it has not been written up methodologically and development work was only now completed, including documentation, a test-dataset, step-by-step tutorial and a largely improved user-interface.

In this manuscript, we apply *Lead Group* to investigate a previously published retrospective cohort of 51 patients suffering from Parkinson’s disease (PD) that underwent STN-DBS surgery. Electrodes of the whole group are visualized both in two- and three-dimensional views. Different types of variable mappings are presented and novel types of connectome derived approaches are investigated. We present concise results and release this dataset in form of a *Lead group* project. A step-by-step tutorial that allows for reproduction of core result figures presented here is included as supplementary material.

**Figure 1:**
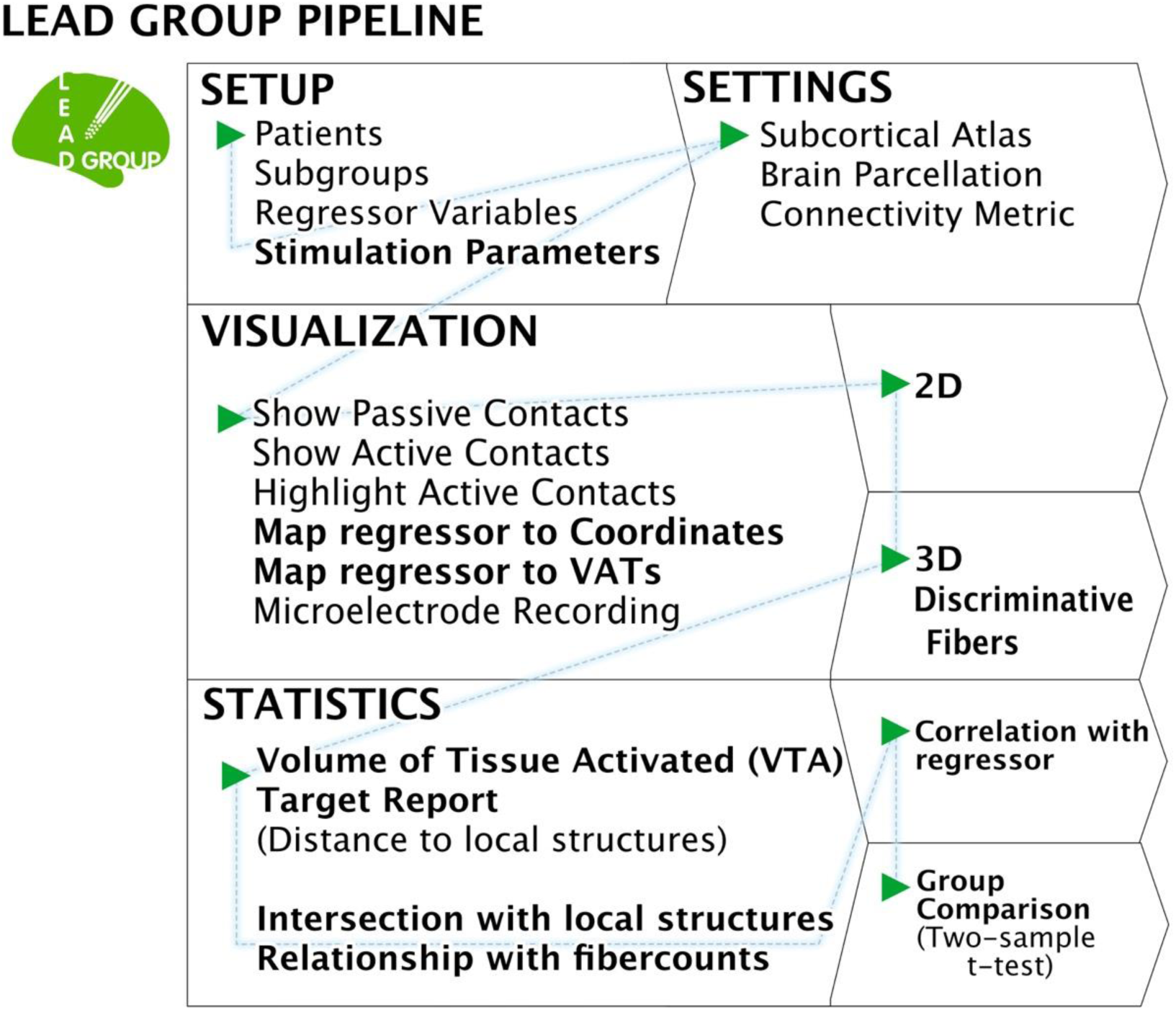
Lead Group Pipeline. The main features and options of the toolbox are shown, including the setup of a group study and settings, which can be chosen for the following processing steps: The visualization of all electrodes in 2D and 3D as well as statistical analyses in relation to either local or connected structures. Those stages highlighted in bold text are described in more detail in the paper.

## Methods and Material

### Patient Cohort

To illustrate the multiple analysis pipelines and visualization options available within *Lead Group*, we included data from a retrospective cohort, which has been described in detail elsewhere (Horn et al., 2019a). Briefly, fifty-one patients implanted with 2 quadripolar DBS electrodes (Model 3389; Medtronic, Minneapolis, MN) to bilateral STN to treat Parkinson’s disease (PD) were included. Patients underwent DBS surgery at Charité – Universitätsmedizin Berlin. DBS response was defined as the percentage improvement along the Unified Parkinson’s Disease Rating Scale (UPDRS)-III motor score ON vs. OFF, which was assessed within an interval of 12-24 months after surgery. Clinical ratings took place after a washout from dopaminergic medication of more than 12 hours. Imaging data of all patients consisted of multispectral preoperative MRI (T1 and T2 weighted) sequences scanned at 1.5T, axial, coronal and sagittal postoperative T2 sequences for 45 patients or postoperative CT scans for the remaining 6 patients. This study was approved by the local ethics committee of the Charité, University Medicine Berlin (master vote EA2/186/18).

### Electrode Localization

Before patient folders may be imported and further processed in *Lead Group*, it is necessary to localize electrodes for each patient using Lead-DBS (which is optimized for single-patient use, bulk-processing with parallel computing, or job submission systems on compute clusters). Localizations were carried out using default parameters of the Lead-DBS v.2 pipeline (Horn et al., 2019a). Briefly, linear co-registration of postoperative images to preoperative MRI scans were performed using a linear transform solved using Advanced Normalization Tools (ANTs; http://stnava.github.io/ANTs/; (Avants et al., 2008)). If needed, these linear within-patient transforms were manually refined using 3D Slicer (www.slicer.org). Preoperative scans were multispectrally normalized into MNI (ICBM 2009b NLIN Asym; (Fonov et al., 2009)) space using ANTs and the “Effective: low variance” protocol with subcortical refinement implemented in Lead-DBS. This normalization scheme was top performer in a recent exhaustive evaluation for registrations of subcortical structures such as the STN and GPi (Ewert et al., 2019b).

### Lead Group Setup and Visualization

To set up the group analysis in *Lead group*, all 51 patients were selected and their percentage improvement on the UPDRS-III motor score entered as a variable into the *Lead Group* GUI (see section S1.1 in the walkthrough tutorial appended within the supplementary material). Some analyses offered by Lead Group are parametric in nature, i.e. could directly take advantage of the continuous variable *%-UPDRS-III improvement*. However, others are meant to compare groups (e.g. to analyze differences between hospitals or compare top vs. poor responders). To demonstrate these analysis streams, as well, a median split was applied to assign a variable separating good (25) from poor (23) responders, with 3 patients lying exactly on the median score of 44 % (not assigned by the variable). This step arbitrarily separated the group in two subgroups of similar size. Stimulation parameters, i.e. active contacts and amplitudes, were specified for each individual patient. Volumes of tissue activated (VTA, representing a rough approximation of the surrounding tissue modulated by DBS) were calculated using a finite element method (FEM) approach (Horn et al., 2019a, 2017c). As alternatives, three other heuristic models are implemented in Lead-DBS (Dembek et al., 2017; Kuncel et al., 2008; Mädler and Coenen, 2012) which were not used in the present manuscript.

To graphically illustrate the electrode placement in relation to respective clinical outcome following DBS, active contacts of all patients from the two groups (upper vs. lower half in clinical improvement) were visualized, both in two-dimensional and three-dimensional space (sections S1.2 - S1.4 in the walkthrough tutorial). A 100-micron T1 scan of an ex vivo human brain, acquired on a 7 Tesla MRI scanner with a scan time of 100 hours (Edlow et al., 2019) served as a background template throughout this paper and the definition of STN boundaries was informed by the DISTAL atlas (Ewert et al., 2018a).

In a next step, four different implemented options that map variables (such as %- UPDRS-III improvement in the example case) to anatomical space were applied. First, all active contacts were visualized as a point cloud and colored by the variable intensity (section S1.6 in the walkthrough tutorial). Second, the improvement variable was mapped to an equidistant point grid after solving a scattered interpolant based on the original values and active coordinates (section S1.7 in the walkthrough tutorial). Third, this equidistant grid was visualized as an isovolume, i.e. the grid was thresholded and visualized as a 3D-surface (section S1.8 in the walkthrough tutorial). These first three options mapped clinical improvement to the *active contact coordinates*. A fourth option is available that instead mapped these values to *VTAs* (section S1.9 in the walkthrough tutorial). Here, each binary VTA was weighted by its corresponding improvement value and for each voxel a T-value was estimated. The resulting volume was thresholded based on visual inspection at an arbitrary T-value (10.56). This value was visually chosen to obtain a “sweet-spot” with small anatomical extent.

### Connectomic analyses

Seeding from VTAs, estimates of structural connectivity to other brain areas can be computed in *Lead group*. To do so, patients’ individual diffusion weighted images or a normative connectome may be used. Within Lead-DBS, four structural group connectomes are currently available and can be chosen depending on the cohort of study. For the present work, a PD-specific connectome was used, which was obtained from 85 patients included in the Parkinson’s Progression Markers Initiative (PPMI; www.ppmi-info.org) database (Marek et al., 2011). This processed connectome has been used in prior studies (Ewert et al., 2018a; Horn et al., 2019a, 2019b, 2017a, 2017c; Irmen et al., 2019b) and processing details are reported elsewhere (Ewert et al., 2018a).

For the present analysis, we selected fibers of the connectome that traversed through the VTA and terminated in distinct regions of a publicly available parcellation of the sensorimotor cortex, the Human Motor Area Template (HMAT; Mayka et al., 2006), which contains regions defining S1, M1, supplementary and presupplementary motor area (SMA/preSMA), dorsal and ventral premotor cortex (PMd/PMv; section S1.12 in the walkthrough tutorial).

A second connectomic analysis stream implemented into *Lead group* works on a fiber level in a mass-univariate fashion (section S1.13 in the walkthrough tutorial). For each tract of a group connectome, improvements associated with VTAs that are connected to the tract are compared to the improvement values of VTAs not connected to the tract in two-sample t-tests. The method is referred to as “discriminative fibertract analysis” in the software and was introduced in (Baldermann et al., 2019). By running the aforementioned test for every tract, each receives a predictive value in form of a T-score that can be positive or negative. Tracts are then color-coded by their T-score for visualization. Fibers mapped in red are predominantly connected to VTAs that were associated with better treatment response. The opposite would account for “negative fibers”. These are not shown given the large overall improvement of the cohort. Also, based on clinical and pathophysiological knowledge, it is unreasonable to hypothesize that DBS with poor placement would contribute to worsening of motor symptoms above and beyond side-effects which are not covered by the UPDRS and not largely present in long-term stimulation-settings as the ones studied here.

### Statistical Analysis

While the aforementioned processing steps mainly serve to visually describe DBS effects with regard to their anatomical sites, *Lead group* further provides specific statistical tests and ways to export metrics to run more elaborate statistical analyses in different software. For instance, it is straight-forward to export intersection volumes between VTAs and a specific anatomical atlas structure (such as the STN) or correlate these with clinical improvement values directly within *Lead group*. For the purpose of this study, intersections between each patient’s (bilateral) VTA and the (bilateral) STN atlas volume were correlated with the clinical outcome variable by conducting a Spearman’s rank-correlation. Random permutation (× 5000) was conducted to obtain P-values. Similarly, the values of the top and bottom responding half of the cohort were compared using a two-sample T-test. The same two types of analyses can be applied to connectivity estimates (e.g. of tract counts connecting VTAs with a cortical region such as the SMA). Similarly, those metrics can instead be exported for further analysis elsewhere.

## Results

### Clinical Improvement

The 51 patients (age 60 ± 7.9; 17 female) improved by 45.4 ± 23.0% on the UPDRS III motor score, from a postoperative baseline of 38.6 ± 12.9 to 21.1 ± 8.8 points. For further demographic details on the cohort, please see (Horn et al., 2019a, 2017c).

### Electrode placement

Active contacts were mapped to anatomical space and visualized in 2D slice views (figure 2). Contacts of the top responding half are shown in light red (and the best-responding patient in dark red) while the bottom half is shown in light blue (with the poorest responder in dark blue). A similar export is shown in 3D in the left panels of figure 3 which separately shows the two medium-split halves of active contacts. The right panel shows 3D electrodes with realistic dimensions instead of point-clouds.

**Figure 2:**
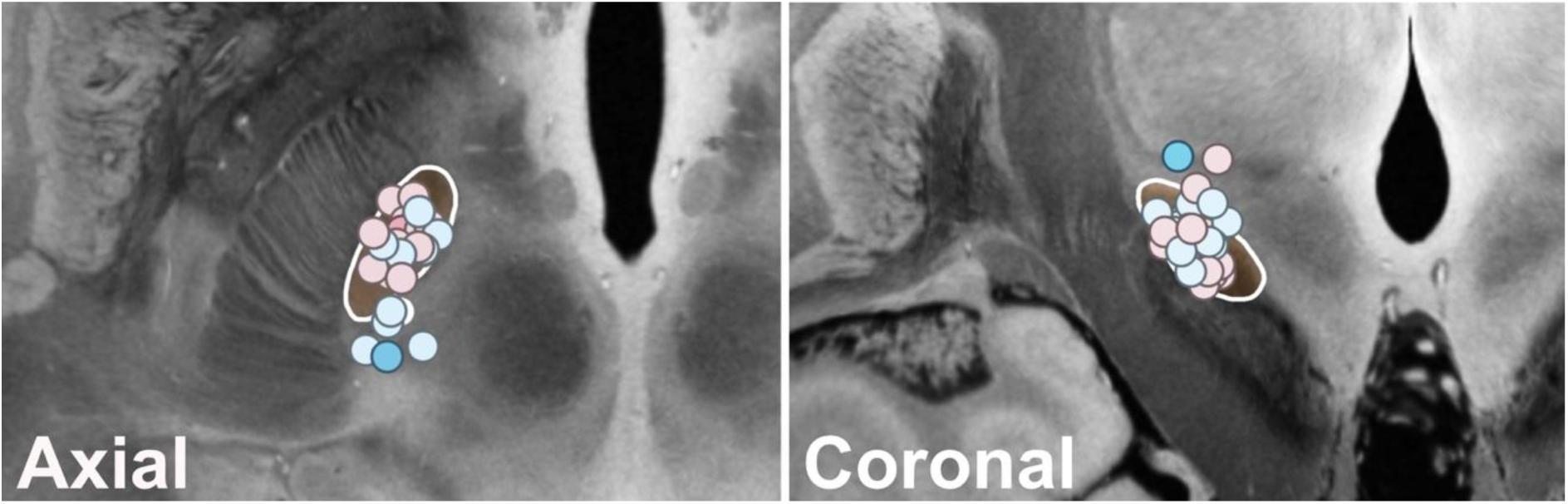
2D Visualization in Lead group. 2D slices of the most dorsal contacts in the left hemisphere (K9). Axial view on the left, coronal view on the right. Light blue and light red colors represent poor vs. good responders, respectively and dark blue vs. dark red circles depict the two patients with the least and most clinical benefit. STN is shown in orange.

**Figure 3:**
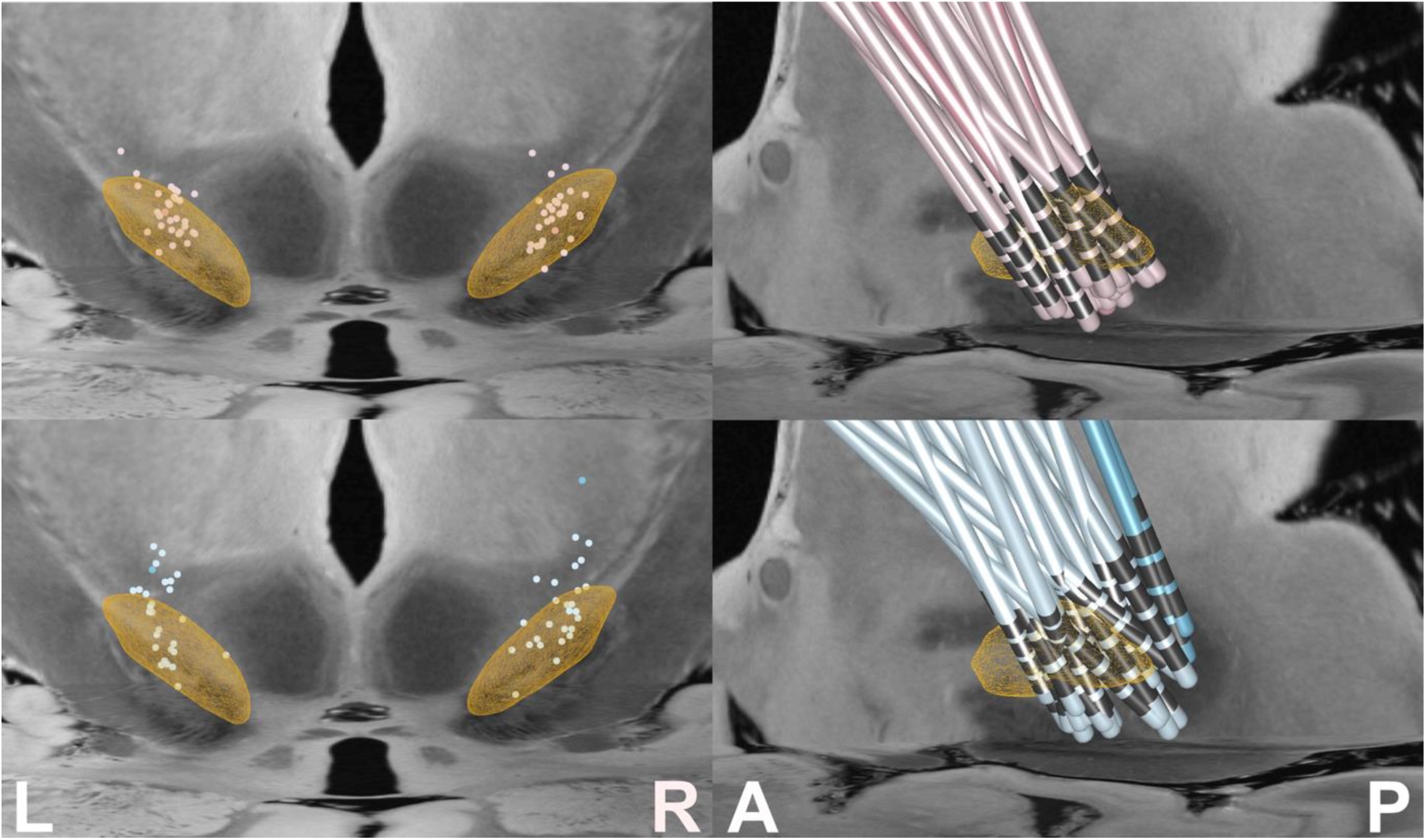
3D visualization in Lead group. *Left side:* All active electrode contacts are shown as Point-Clouds from posterior. *Right side:* Electrodes were mirrored and are shown from the left side. Good and poor responders are presented in the upper and lower row, respectively.

### Electrode position weighted by clinical improvement

After the clinical regressor was mapped to coordinates of active electrode contacts and VTAs, these were visualized in various forms using Lead group in 3D (figure 5) and exported to visualize them in a NIfTI-viewer such as 3D Slicer in 2D (figure 4).

**Figure 4:**
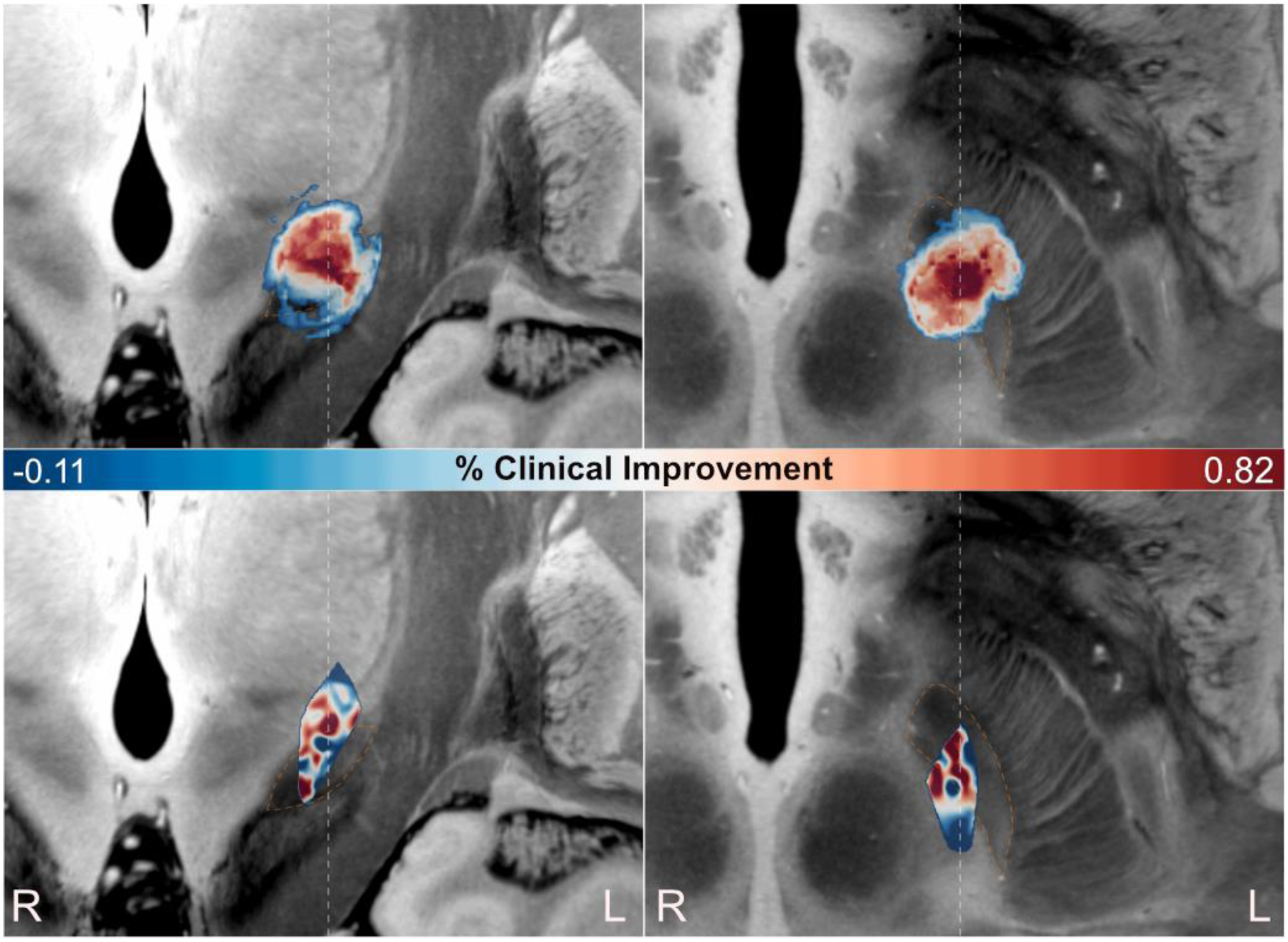
Clinical regressor. *Upper panel:* coordinates of all active contacts, weighted by clinical improvement. *Lower panel:* Volumes of tissue activated of all patients, weighted by clinical improvement. Tmaps, corrected for number of VTAs are presented, ranging from negative (blue) to positive (red) clinical outcome. Coronal view on the left, axial view on the right. STN highlighted with dashed orange line. Both regressions point towards an optimal stimulation site within the more dorsal part of the STN.

**Figure 5:**
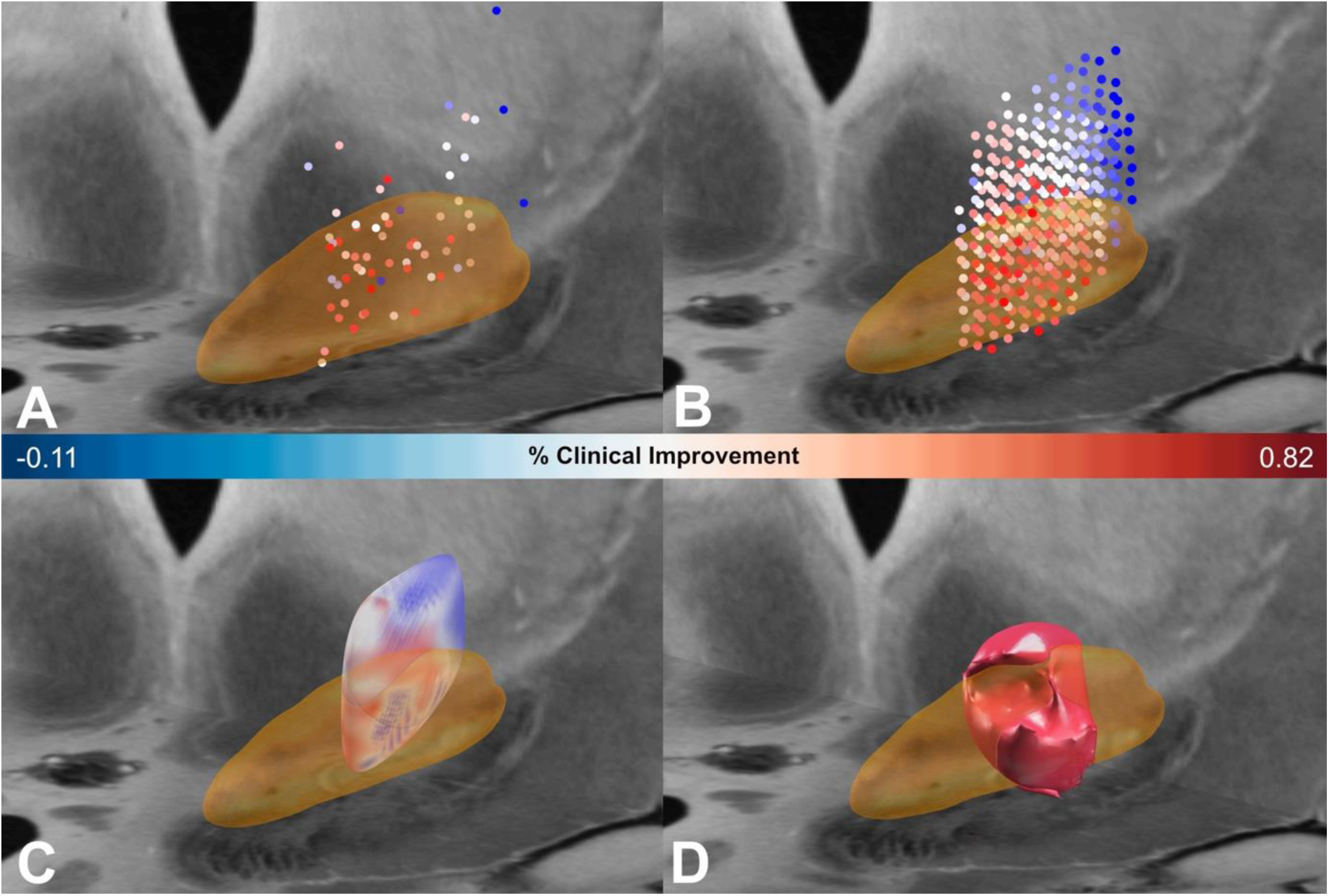
Mapping regressors in Lead group. *Panel A:* All active contacts, shown as point clouds colored by clinical improvement. *Panel B:* regressor is mapped to electrode coordinates as interpolated point mesh. *Panel C:* regressor mapped as isosurface. *Panel D:* VTAs of patients weighted by clinical improvement and thresholded by a T-value of 10.56; Color bar relates to Panels A-C and clinical improvement is guided by percentage UPDRS-III motor score improvement. Left hemisphere is displayed, with STN in orange.

Figure 5A shows the active contacts of both sides that were nonlinearly flipped to the left hemisphere. Contacts were color-coded by clinical improvement values, showing a predominantly better improvement within the STN as opposed to outside of it. This scattered point cloud was then used to fit a scattered interpolant from which an equidistant volumetric grid could be generated. In figure 5B, the raw points of this interpolation grid are visualized, leading to a potentially clearer impression of the spatial extent of the optimal target region. Figure 5C uses a different approach to visualize the same interpolated data grid in showing a 3D isosurface that is color coded by improvement values in surrounding points. By doing so, a point cloud (of active coordinates) is transformed to a volume which is shown as 2D slices in the upper panel of figure 4.

Mentioned methods use active coordinates, while it is also possible to map improvement values to the spatial extent of each VTA. Figure 5D and bottom panel of figure 4 show results of voxelwise T-maps across weighted VTAs.

Independent of method or visualization strategy, these results favor an optimal target within the STN and at anterior level of the red nucleus at its largest extent. This confirms priorly published articles that came to the same conclusion (Akram et al., 2017; Bot et al., 2018; Horn et al., 2019a).

### Distance to target

Above approaches aimed at defining optimal targets in a data-driven fashion, i.e. by weighting electrode coordinates or VTAs with clinical improvement scores and aggregating those values on a group level. Alternative research questions could aim at validating *known* targets or coordinates. For instance, if an optimal target coordinate was reported in the literature, an aim could be to validate the target using a novel cohort (Horn et al., 2019a). Here, we explore this option by calculating distances between each target coordinate and the atlas definition of the STN. To do so, a “target report” can be generated within *Lead group* (see section S1.10 in the walkthrough tutorial), which calculates the distance in mm of each electrode contact to the closest voxel of the chosen atlas structure. In addition, a threshold can be selected so that a binary table will be provided, indicating whether or not a contact resides inside or outside of the anatomical structure. Of note, this “structure” can be defined as any map defined in NIfTI format in standard space, i.e. it does not necessarily need to be a brain nucleus like the STN in our example. In figure 6 these distances are presented for the best (82% improvement) and the poorest (−11% improvement) responders in synopsis with their electrode reconstructions. Not surprisingly, their proximity to the target differs largely and the active contact of the best responder was placed inside the STN.

**Figure 6:**
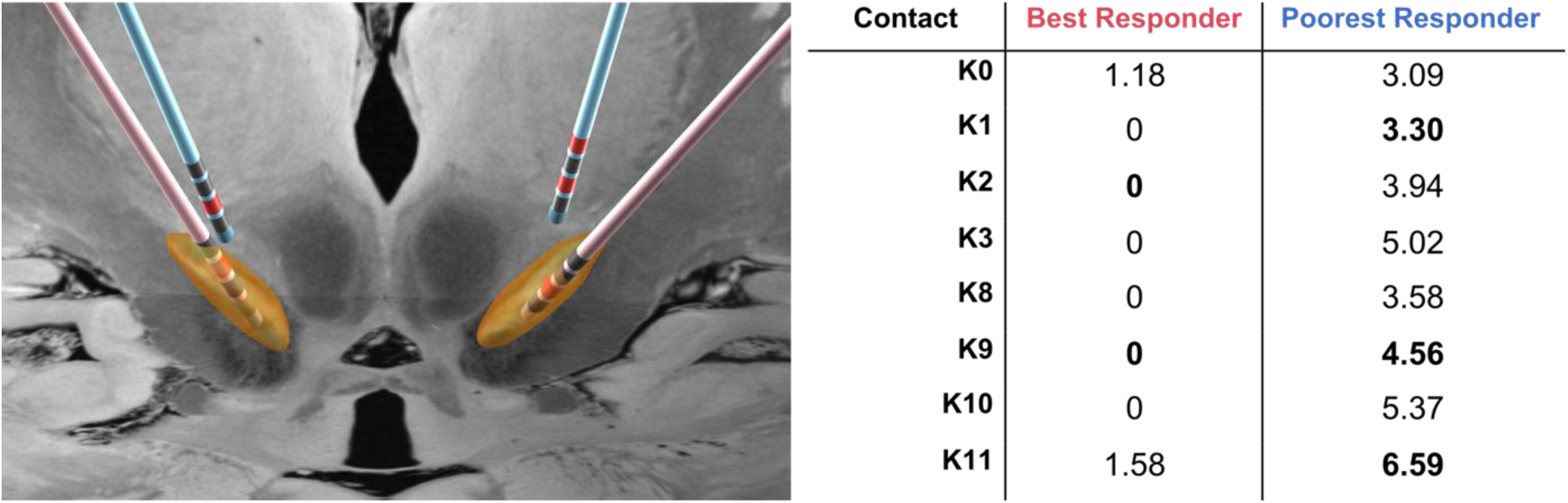
“Target report” feature showing distance from contact center to its closest STN voxel. For each active contact (highlighted in red), the distance from the contact center to the STN is shown for patients with best (red) and worst (blue) clinical outcomes.

### Intersection with local structures

Instead of calculating absolute electrode *distances*, a similar approach is to calculate the *intersections* between each VTA and an anatomical atlas structure (see section S1.11 in walkthrough tutorial). These intersecting volumes may be either used in statistical tests in *Lead group*, or exported for further analysis. In our example, we hypothesized, that clinical improvement would correlate positively with the volume of STN intersection and accordingly, that the group of top half responders, as initially assigned by a median split, would intersect with the STN to a significantly higher degree than the group of bottom half responders (figure 7).

**Figure 7:**
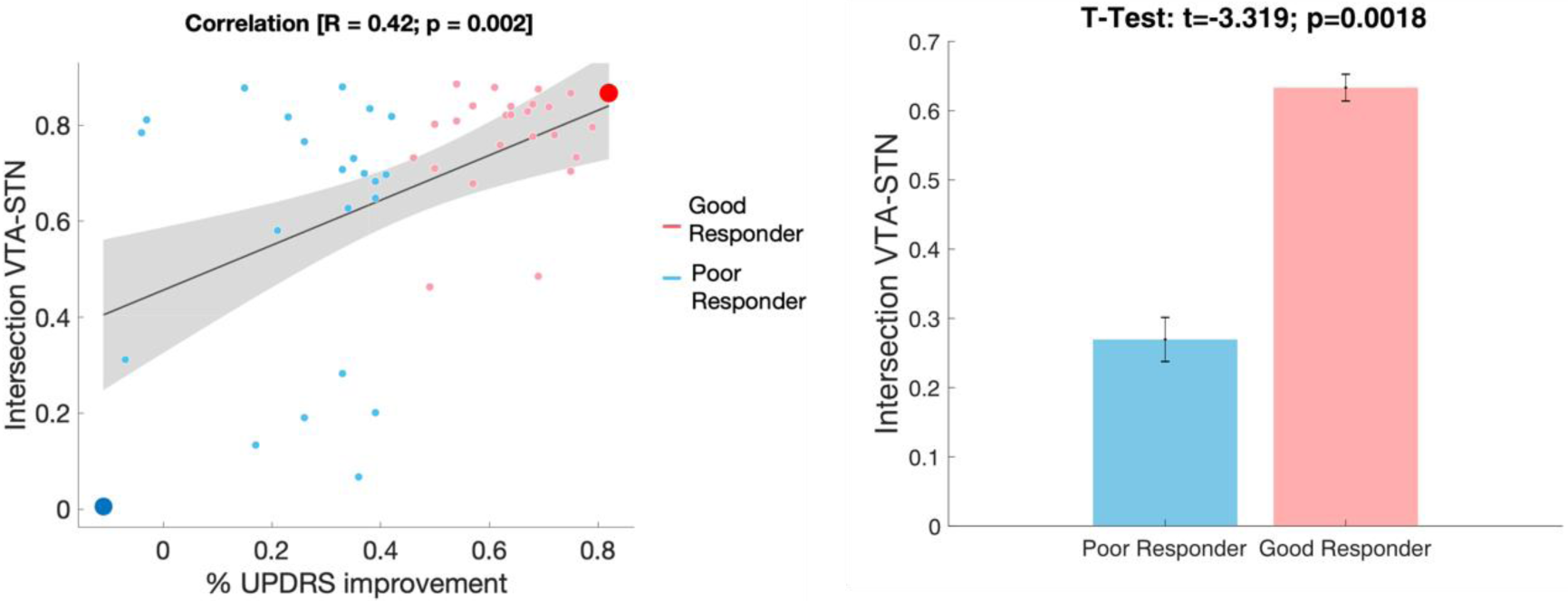
VTA intersection with the STN explains clinical improvement. *Left side:* Spearman’s rank-correlation between clinical improvement and intersections between VTA and the STN. Patients with best vs. poorest responses (figure 6) are highlighted with large circles. *Right side:* Patients were arbitrarily median-split into two groups, based on their percentage improvement on the UPDRS-III score. The group of better responders (red) showed significantly higher VTA overlap with the STN.

### Fibercounts to connected structures

Based on the PPMI 85 connectome, fibers traversing through each VTA were isolated and the ones terminating in each cortical region defined by the HMAT parcellation were counted using *Lead group.* These numbers were correlated with clinical improvement across the group. As can be seen in figure 8, the number of fibers reaching preSMA (R = 0.29) and PMv (R = 0.3) could explain part of the variance in clinical improvements. In turn, less fibercounts connecting VTAs to M1 or S1 were associated with better %UPDRS-III improvement (R = −0.4 and R = −0.46). Of these results, only the latter two would remain significant after applying Bonferroni correction to p-values.

**Figure 8:**
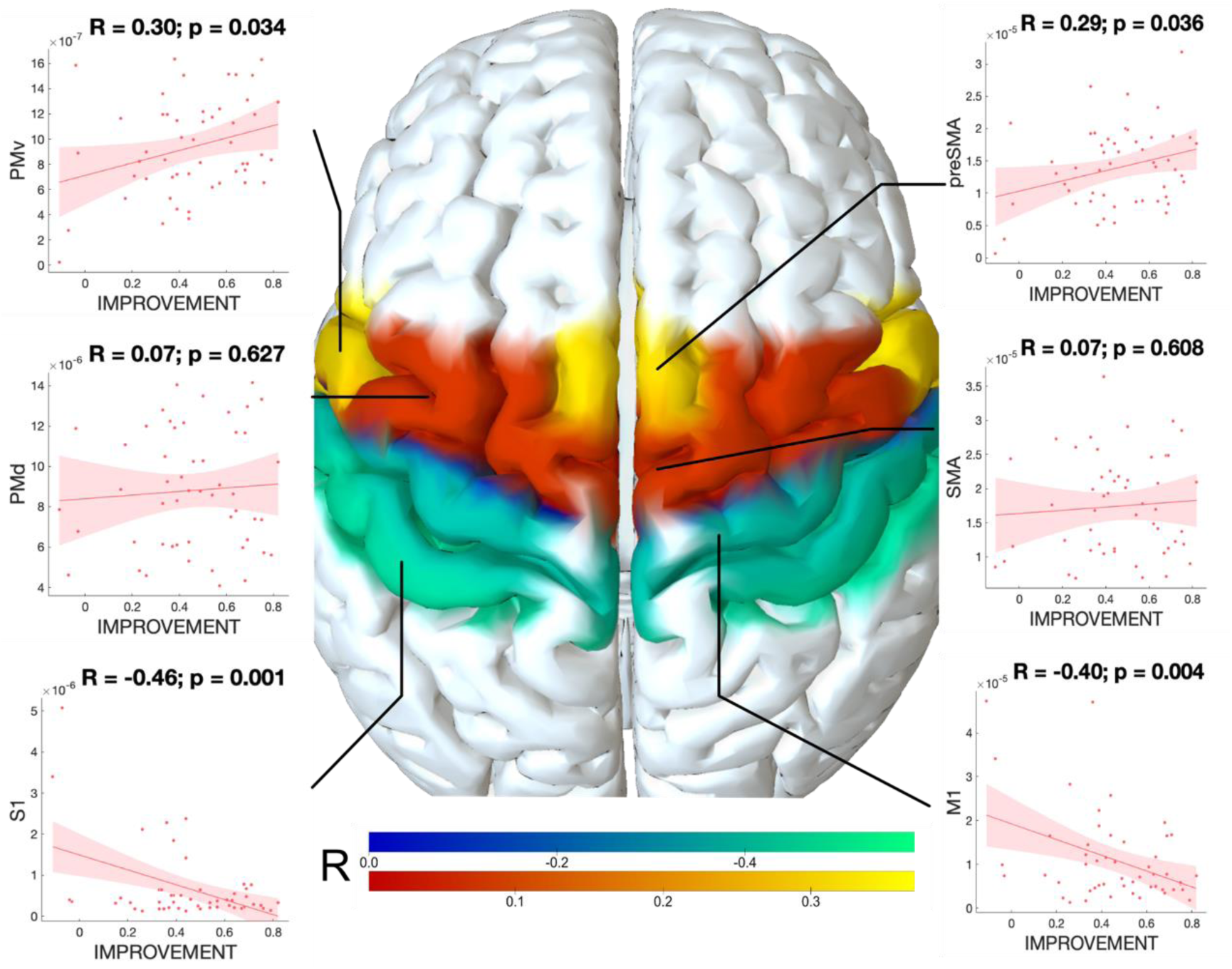
Correlation with connectivity to cortical structures. The six structures of the HMAT parcellation, colored by the R value, resulting from Spearman’s rank-correlations between fibercounts to these areas and clinical improvement.

### Discriminative fibers

The above analysis aimed at identifying cortical regions to which connections were positively or negatively associated with clinical improvements. An additional analysis stream available within *Lead group* could complement this concept by identifying specific *tracts* that were associated with clinical improvement. To do so, mass-univariate T-tests were conducted for each tract of the normative PD connectome between improvement values of connected vs. unconnected VTAs. Fibers were then color-coded by their T-value. The top 10% of the fibers are shown in figures 9A and 9B. This analysis revealed that the most positively associated tracts of the connectome traversed within the internal capsule and seemed to branch off to the STN in a similar fashion and at a similar location as the hyperdirect pathway when revealed by single-axon tracing in the macaque (e.g. compare to figs. 2-4 in Coudé et al., 2018a or fig. 4 panel B subpanel v in Petersen et al., 2019). Crucially, tracts that passed by the STN (i.e. fibers of the cortico-spinal tract) received lower T-values and are not shown due to thresholding applied to results. This is remarkable, since the hyper-direct pathway (i.e. corticospinal tract axons that branch off collaterals to the STN) is considered hard if not impossible to differentiate from the corticospinal tract based on diffusion-based tractography (Petersen et al., 2019a). To address this and similar shortcomings of diffusion tractography, Petersen et al. have recently genuinely applied holographic manual reconstructions to define an atlas that defines hyperdirect pathways to the STN (Petersen et al., 2019a). The lower limb M1 portion of this atlas is shown in direct synopsis with our results (figure 9 C).

**Figure 9:**
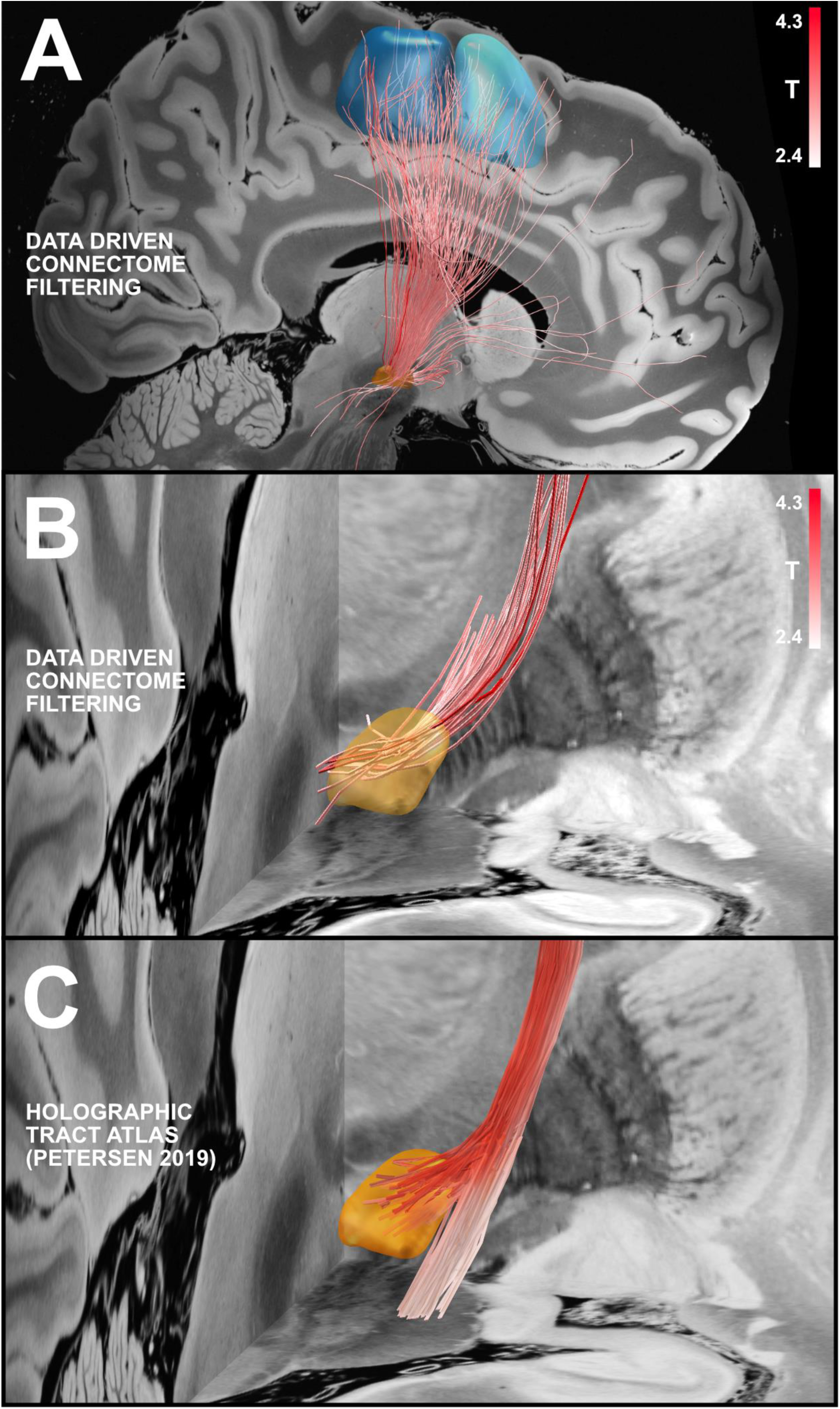
Discriminative fibers. Tracts that are positively associated with clinical improvement are colored by their T-value (in panels A & B). Fibers with strongest discriminative value pass from the STN via the internal capsule to the motor cortex. SMA (dark blue) and preSMA (light blue) which were informed by the HMAT atlas (A). Within the STN (B), the most positively significant fibers (shown from posterior) traverse within the internal capsule and seem to branch off to the STN in a similar fashion as the hyperdirect pathway (e.g. compare to figs. 2-4 in Coudé et al., 2018). Moreover, these tracts project to the dorsal part of the STN at the same location that peaked when mapping improvements to active coordinates or VTAs (figure 4 and 5). Please note that the hyperdirect pathway is implemented by axon collaterals of axons that pass by the STN as can be seen in direct synopsis with an atlas of the hyperdirect pathway (Petersen et al., 2019); C) in which the collaterals are shown in red while the passing axons were faded to white.

## Discussion

Advanced neuroimaging methods are, without any doubt, inevitable to advance Deep Brain Stimulation. Not only can they assist surgical targeting, but also post-operative analyses of DBS effects and to gain added knowledge about functional neuroanatomy. Here, we present a freely-available open-source toolbox that was designed to carry out neuroimaging-based DBS research on a group level. Although our own development focused on patients undergoing DBS with chronically-implanted electrodes, *Lead group* was recently used to localize stereotactic intracranial EEG (iEEG) electrodes as well (Toth et al., 2019).

To demonstrate the functionality of *Lead group*, we analyze a retrospective cohort of 51 PD patients that underwent DBS surgery to the STN. This example application of *Lead group* led to results that were mostly confirmatory in nature but give a comprehensive picture of optimal DBS placement in the STN to treat PD. Specifically, our analysis confirms that an optimal DBS target resided within the dorsolateral STN and suggests that the hyperdirect pathway could play a role in mediating treatment effects.

### Neuroimaging and DBS

The use of neuroimaging in the field of DBS has a long-standing history with x-ray and myelography applications as early examples. With the rise of MRI and CT modalities, it became standard to include neuroimaging data into the clinical procedure. For instance, at around 2000, Hariz and colleagues published a paper that reported use of T2-weighted MRI for pre- and postoperative imaging in a time at which most centers still used indirect targeting on T1-volumes or ventriculography (Hariz et al., 2003).

The rise of MRI and CT made it possible in numerous works to report on electrode placement in the individual patient (e.g. Krause et al., 2015; Laxton et al., 2010b for examples). However, while screenshots of postoperative imaging serve the purpose of confirming electrode placements well, they do not make them comparable to other patients and centers, in a standardized way.

Instead, when using normalized and model-based electrode reconstructions in a standardized space, electrode placement may be characterized in a more objective fashion. This concept has been used in early work that characterized electrodes in standardized imaging-based spaces (e.g. Butson et al., 2011; Eisenstein et al., 2014; Frankemolle et al., 2010; Maks et al., 2009; Nowinski et al., 2005). While the neuroimaging field (mostly driven by fMRI literature) largely converged on the MNI space, most of the functional neurosurgery literature expressed standardized coordinates in coordinates relative to the anterior and posterior commissure (AC/PC) (Horn et al., 2017a). This approach has strong limitations because it does not take patient-specific anatomical variability into account (Horn et al., 2017a; Nestor et al., 2014). The Mayberg group may have been among the first to clinically adopt use of the MNI space for DBS localizations, potentially because surgical targets for depression resided more distant to the AC/PC (leading to larger errors) and because modern imaging modalities like fMRI, diffusion MRI and PET formed elementary components of their pioneering research (Choi et al., 2015; Riva-Posse et al., 2017, 2014).

### A tool to shift DBS imaging research to a group level

Despite aforementioned examples that showed the promise of group localizations, an open-source software capable of performing these analyses has not been developed (while a commercial solution is available in form of the CranialCloud software (D’Haese et al., 2015)). Lead-DBS and *Lead group* were specifically designed to perform group analyses in the field of DBS imaging and have been applied in over 120 peer-reviewed articles (see https://www.lead-dbs.org/about/publications/). Examples that utilized group statistics span across different diseases like PD (Bouthour et al., 2019; Horn et al., 2019c), essential tremor (Al-Fatly et al., 2019; Kroneberg et al., 2019), Dystonia (Neumann et al., 2017), Meige syndrome (Yao et al., 2019), OCD (Baldermann et al., 2019; Huys et al., 2019), epilepsy (Middlebrooks et al., 2018; Wang et al., 2019), Tourette’s Syndrome (W. J. Neumann et al., 2018) or refractory thalamic pain syndrome (Levi et al., 2019). Using Lead-DBS, a sweet spot for STN-DBS in PD was defined and used to predict improvement of motor symptoms in out-of-sample data (Dembek et al., 2019a). Different subregions of the STN were associated with different non-motor outcomes in second study, by exploring local DBS effects (Petry-Schmelzer et al., 2019). Side-effects, such as hyperhidrosis (Yang et al., 2019) or ataxia and dysarthria (Al-Fatly et al., 2019) have been mapped to anatomy using a prototype of *Lead group*. In a sample of epilepsy patients implanted for DBS to the anterior nucleus of the thalamus (ANT), functional connectivity seeding from the VTAs was analyzed with respect to clinical response, using a normative connectome (Middlebrooks et al., 2018). A different study applied *Lead group* to investigate a novel parietal surgical approach for ANT-DBS in epilepsy (Wang et al., 2019). In another recent study, resting-state functional MRI was acquired in PD patients while DBS was simultaneously switched on and off (Horn et al., 2019c). Here, DBS was able to shift functional connectivity profiles towards the ones observed in healthy controls. Focusing on a different analysis path, electrophysiological measures recorded from the LFP-signal of STN-DBS electrodes were mapped to subcortical anatomy of the human brain (Horn et al., 2017b). This confirmed that elevated beta-power is predominantly expressed within the sensorimotor functional zone of the STN, a finding that was reproduced and extended by two different teams again by the use of a *Lead group* prototype (Geng et al., 2018; van Wijk et al., 2017). The concept is now referred to as *subcortical electrophysiology mapping* and the involved research questions shaped functionality and continued development of *Lead group* while being applied in further studies (Lofredi et al., 2018; Neumann et al., 2017; Tinkhauser et al., 2019).

The ability to nonlinearly map electrodes of patient cohorts into a comparable and well-defined space also led to the possibility of exploring subtle differences in their brain connectivity profiles. After pioneering work that applied commonly available pipelines from the neuroimaging field or commercial tools (Akram et al., 2018; Vanegas-Arroyave et al., 2016), *Lead group* was further improved to perform such analyses, as well (Horn et al., 2017a). Since then, the tool has empowered research that explored optimal connectivity profiles in PD (Horn et al., 2017c), OCD (Baldermann et al., 2019) and Essential Tremor (Al-Fatly et al., 2019).

Extending this line of research, it was used to relate connectivity profiles seeding from DBS electrodes to *behavioral* instead of clinical changes induced by DBS. For instance, Neumann et al. showed that specific connections of the electrodes would lead to changes in movement velocity vs. reaction times in a motor task(W.-J. Neumann et al., 2018). De Almeida showed that functional connectivity between STN-DBS electrodes and a specific site in the ipsilateral cerebellum was associated with partly restoring motor learning in PD patients (de Almeida Marcelino et al., 2019). Finally, focusing on localized instead of connectivity-mediated effects, Irmen et al. showed that modulating specific subregions of the STN could restore risk-taking behavior to a level observed in healthy controls (Irmen et al., 2019a).

### Optimal placement in STN-DBS for treatment of PD

While above results demonstrate general benefits of a DBS imaging group level tool, results obtained from the example cohort in the present manuscript should be set into scientific context, as well.

In summary, our results confirm previous findings, that the stimulation of the STN, and more precisely its dorsolateral part is linked to UPDRS-III improvement (e.g. Caire et al., 2013). As mentioned above, this finding is not novel. In fact, recent studies that were carried out by four centers, each using three different methodologies, converged on almost the exact same optimal target coordinate (Akram et al., 2017; Bot et al., 2018; Horn et al., 2019a; Nguyen et al., 2019; for a review see Horn, 2019). One of those studies used commercial software (Akram et al., 2017), one a method that directly builds upon the imaging data in native patient space (Bot et al., 2018) and the third and fourth used *Lead group* (Horn et al., 2019a; Nguyen et al., 2019). Moreover, the two latter studies were able to significantly explain clinical improvement by measuring distance from each DBS electrode to the optimal coordinate. This illustrates that the precision of DBS imaging research has evolved to become useful, after a list of earlier studies had partly shown conflicting results (Horn, 2019).

Our results further showed that treatment success was positively associated with structural connectivity to preSMA and PMv. Again, this has been shown before (Akram et al., 2017; Horn et al., 2017c; Vanegas-Arroyave et al., 2016). Here, we elaborate on this finding by isolating a specific fiber bundle that seems to be associated with clinical improvement. Interestingly, this analysis revealed that fiber-tracts that connect the motor cortex with the STN are most associated with good clinical improvement while tracts of passage (i.e. the corticospinal tract, CST) are less so (figure 9). This result may be surprising, since separating the hyperdirect pathway from the CST is considered difficult if not impossible by means of dMRI based tractography (Petersen et al., 2019a).

In the brain, the hyperdirect pathway is implemented in form of axon-collaterals branching off from axons traversing from the cortex inside the internal capsule (Coudé et al., 2018b; Gunalan et al., 2017; McIntyre and Hahn, 2010; Nambu et al., 2002; Petersen et al., 2019b). Reconstructing such *branching* fibers is not possible in available tractography software packages, for obvious reasons of an already high false-positive rate (Maier-Hein et al., 2017). Still, in direct comparison with single-axon tracing data in the macaque, the anatomical course of the tract isolated here (figure 9B) starkly resembles the path of the axon-collaterals of the hyperdirect pathway (see tract atlas in fig. 9C or figs. 2-4 in Coudé et al., 2018).

### Limitations

Mapping DBS electrodes to a group template inherently comes with a loss of precision and a multitude of related problems and limitations. Already beginning within the patient’s own space, brain shift introduces non-linear displacements between postoperative and preoperative scans, favoring the use of postoperative MRI, which makes it possible to directly visualize both the electrode and target structure in the same space (Hariz et al., 2003). This problem can partly be overcome by applying brain shift correction (Horn et al., 2019a) as applied here, but a residual error should be assumed to remain, especially in patients with large pneumocephalus volumes (Lee et al., 2010). Errors in localizing DBS electrodes themselves may potentially introduce a greater source of bias than could be assumed. To this end, Lead-DBS has incorporated a phantom-validated algorithm for automatic localizations based on postoperative CT volumes (Husch et al., 2018) as well as a multitude of manual refinement and control views. Still, empirical data on observer-dependencies or inter-rater errors of electrode localizations with Lead-DBS are lacking (such studies are currently underway). Nonlinearly warping electrodes into template space introduces an additional bias that cannot be completely overcome. Based on experience when visualizing the same patient in native and MNI space and comparisons with results from other tools, we are confident that localizations in MNI space are meaningful, since they have been used to explain (Horn et al., 2019a; Joutsa et al., 2018; Yao et al., 2019) or even predict (Al-Fatly et al., 2019; Baldermann et al., 2019; Horn et al., 2017d) clinical improvements in a number of studies. Still, the amount of error introduced by nonlinear registrations remains unclear. Some validation of electrode localizations in standard space were made using electrophysiology by various groups (Horn et al., 2017b; Neumann et al., 2017; Nowacki et al., 2018; Tinkhauser et al., 2019; van Wijk et al., 2017). For instance, agreement between electrophysiology-defined and atlas-defined boundaries in standard space showed high agreement across 303 microelectrode recording sites when localizing recording-sites using Lead-DBS (Rappel et al., 2019). Still, even if reconstructions in MNI space are meaningful, we must assume a slight bias since by definition, nonlinear registrations will introduce inaccuracies. Of note, the exact same bias applies when instead warping atlas structures from template to native space (at least when applying diffeomorphic registration strategies which are standard in the field (Avants et al., 2008)). Over the last decade, we aimed at minimizing this nonlinear error by introducing various concepts that ranged from multi-step linear transforms (Schönecker et al., 2009) over the introduction of multiple algorithms (Ewert et al., 2018a; Horn et al., 2019a, 2017a, 2017b; Horn and Kühn, 2015) to the current multispectral default method that was recently empirically validated (Ewert et al., 2019b). Further methodological work such as the possibility to manually refine warp-fields has been published and implemented in Lead-DBS / *Lead group* (Edlow et al., 2019) but was not yet applied in the present work. Despite these efforts that may *minimize* bias introduced by warping electrodes to template space, these can likely never be *overcome* completely. Thus, the gold-standard for localizing DBS electrodes will always remain to work in native patient space, when the research questions we ask allow us to do so. A further limitation of Lead-DBS and *Lead group* is the accuracy of the implemented VTA models. While our pipeline includes the only finite element method based VTA model that is available in an open-source pipeline, more accurate models have been described in the literature (Chaturvedi et al., 2013; Gunalan et al., 2017). Lead-DBS uses a VTA model that thresholds the E-Field magnitude as this practice was suggested to yield good approximations by others (Åström et al., 2015). However, recent results by leading groups in this field suggested significant limitations of the approach (Duffley et al., 2019; Gunalan et al., 2017). Moreover, limitations apply to the *concept* of the VTA itself. Specifically, it is impossible to represent the bioelectrical effects of DBS within a binary volume of any form. Such a representation cannot include information about which axons of which orientation, diameter or neurotransmitter types are modulated. While GABAergic neurons deplete by DBS almost immediately, Glutamatergic neurons do not (Amadeus Steiner et al., 2019; Milosevic et al., 2018b) – leading to differential modulation effects (while recent evidence suggests that the DBS effect in the STN for PD is predominantly mediated by GABAergic neurons (Milosevic et al., 2018a)). Furthermore, models do not incorporate microscale anisotropy (while the use of macroscale anisotropy has been explored (Butson et al., 2006)) or heterogeneity of tissue (fluid-filled regions around neurons and axons lead to strong spreads of current (Sriperumbudur et al., 2018)). This inherent limitation of representing DBS effects as a sphere has led to more advanced concepts such as pathway activation models (Gunalan et al., 2017) which have yet to enter the open-source software field to become broadly applicable (such developments are currently underway (Butenko et al., 2019)).

Instead of analyzing DBS effects on an axonal/neuronal level, *Lead group* works on a *voxel-level*, to analyze connectomic DBS effects. Thus, it approaches modeling problems from a neuroimaging perspective – while others have approached it from a cellular modelling perspective. While this strategy has been useful to predict clinical and behavioral effects and it does not require a multitude of biophysical assumptions, it is limited in the potential predictions that it can theoretically make (Petersen et al., 2019a).

A related limitation lies in the use of normative connectomes. While their utility could be shown to predict various clinical symptoms (Al-Fatly et al., 2019; Baldermann et al., 2019; Horn et al., 2017a, 2017c; Joutsa et al., 2018; Weigand et al., 2018), these datasets lack patient-specificity and should be seen as brain atlases of the wiring diagram of the human brain. And while atlases have been used in the field of DBS since the introduction of the Clarke-Horsley apparatus in 1906 (Clarke and Horsley, 2007), insights derived from atlases should be transferred to individual patients cautiously, if at all (Coenen et al., 2019). Moreover, diffusion-weighted imaging based tractography and resting-state functional MRI, the modalities usually used for noninvasive connectivity mapping, are both highly derived modalities that show a multitude of inherent problems and lead to a high number of false-positive (Maier-Hein et al., 2017) and false-negative (Maier-Hein et al., 2017; Petersen et al., 2019a) results.

In sum, limitations in precision – from registrations and localizations in native space, warping bias in template space and limitations of applied biophysical models and connectomes – add up and make DBS imaging research challenging, especially so on a group level. Strategies to reduce bias include i) acquiring high-quality multimodal preoperative MRI data, ii) accurate postoperative imaging, iii) care- and skillful analysis and meticulous data control in every step of the pipeline and iv) cautious interpretation of results. Over the years, we have identified typical pitfalls and included control views, correction tools and mediation strategies to reduce bias wherever possible throughout the Lead-DBS software package.

### Conclusions

We present a novel open-source toolbox that is designed to shift deep brain stimulation imaging to a group-level. We show utility of the toolbox by presenting largely confirmatory results in a cohort of Parkinson’s Disease patients that underwent deep brain stimulation surgery to the subthalamic nucleus. Specifically, we validate an optimal target within this nucleus that has conclusively been described by four independent groups within the past two years. Furthermore, we show connectivity profiles and a specific tractogram that were associated with good clinical outcome in our analyses. Finally, we discuss findings and applications of group-level deep brain stimulation imaging while making aware of a multitude of limitations that apply to this strand of research.

## Data availability statement

The aggregated dataset in form of a Lead-Group project used to create figures and analyses is openly available (https://osf.io/kj456/). Raw data (patient MRI and postoperative CTs) cannot be openly shared because it contains patient information. All code used to analyze data presented in the present manuscript is openly available within the Lead-DBS software suite (https://github.com/netstim/leaddbs; https://www.lead-dbs.org/). A step-by-step instruction manual to reproduce findings is available in the supplementary data of the present manuscript.

## Acknowledgement

This study was supported by the German Research Foundation (DFG grant SPP2041, “Clinical connectomics: a network approach to deep brain stimulation” to AAK as well as Emmy Noether Grant 410169619 to AH) and an FPI Predoctoral Fellowship (BES-2016-079470) to ST.

## Supplementary Material

**Figure S1.**
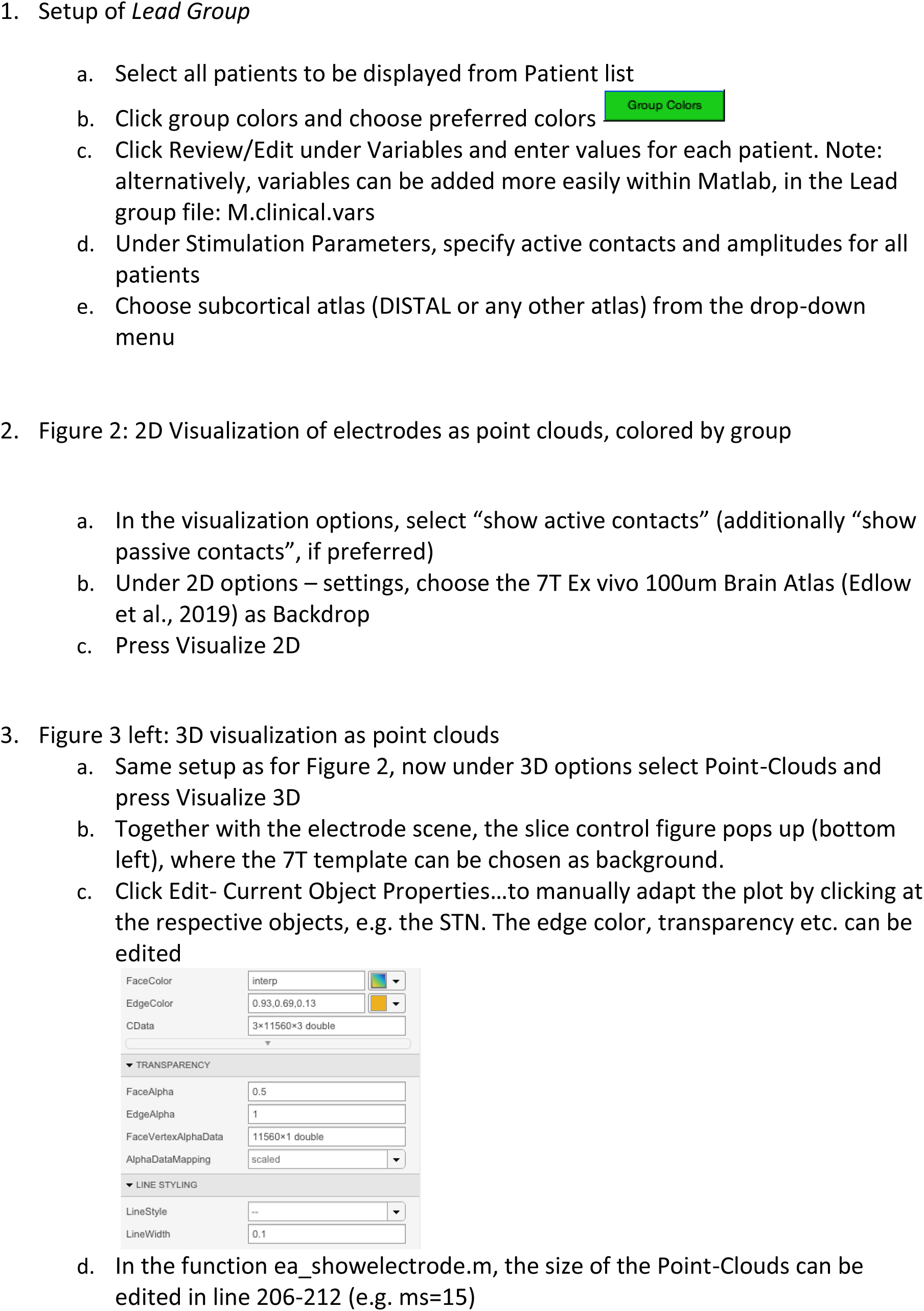
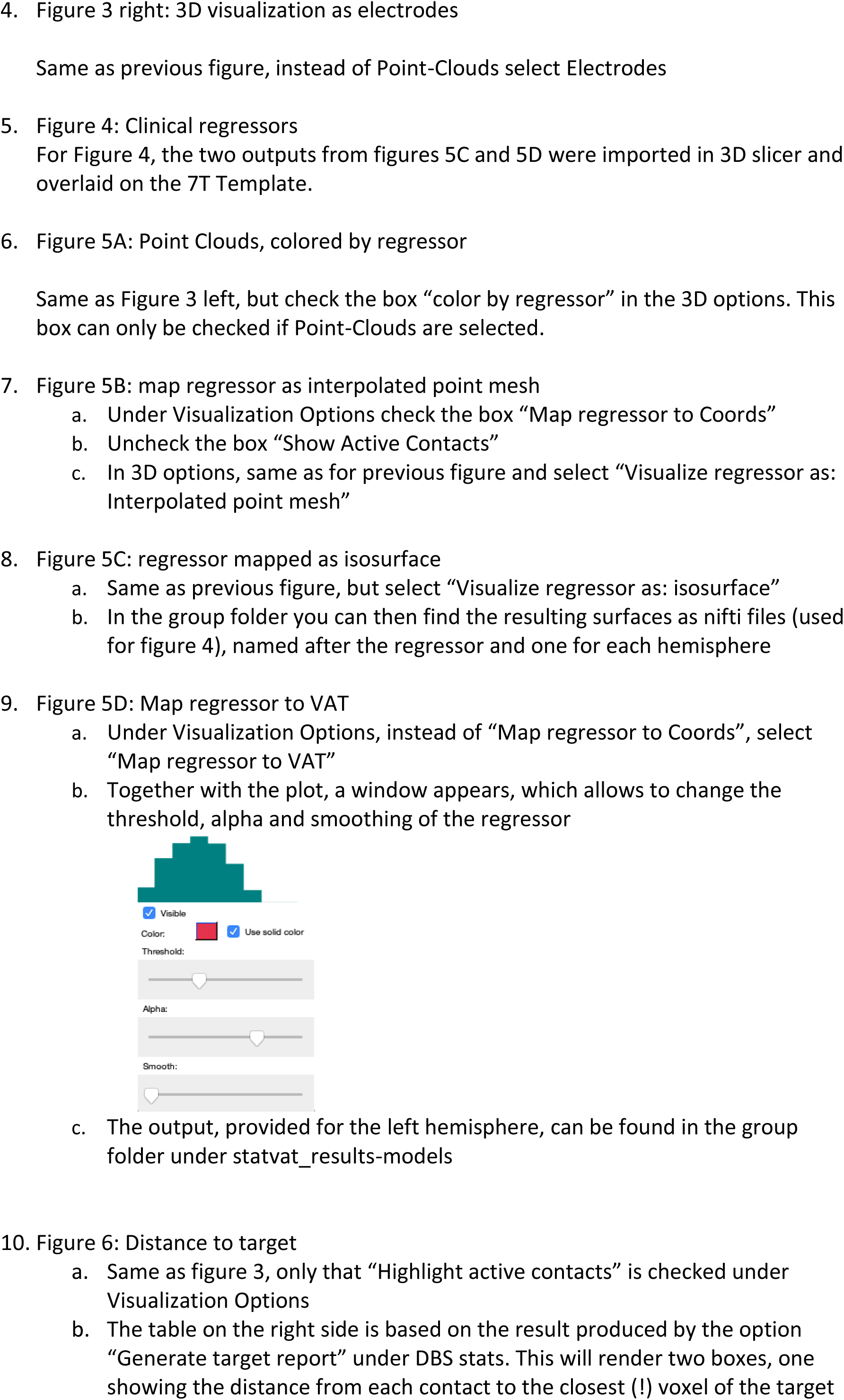
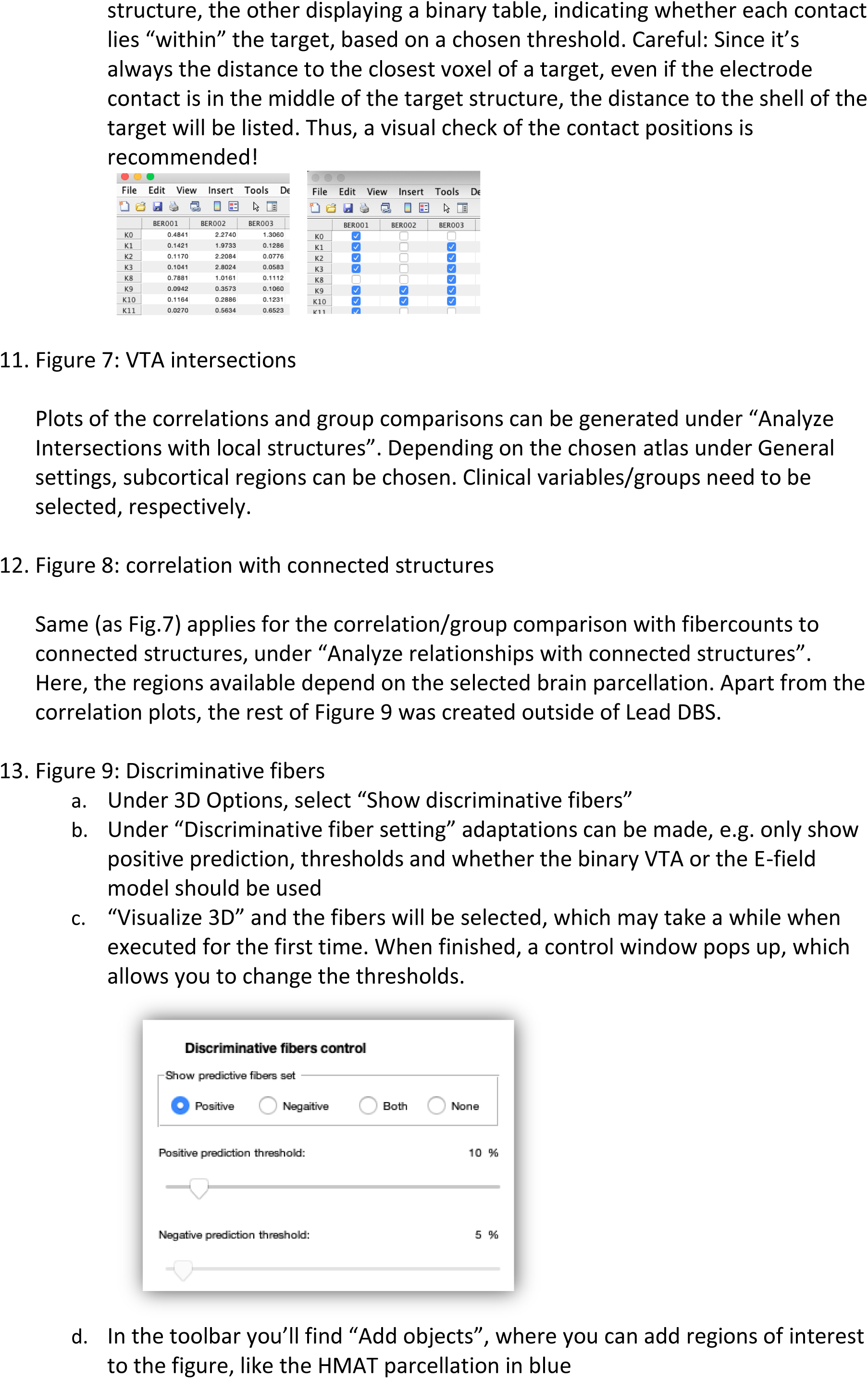
Walkthrough Tutorial.

